# Ubiquitinome analysis of articular cartilage post mechanical injury reveals a differential ubiquitination pattern of a subset of DUBs and proteins linked to the ERAD cellular response

**DOI:** 10.1101/2022.01.19.476879

**Authors:** Nitchakarn Kaokhum, Adán Pinto-Fernández, Mark Wilkinson, Benedikt Kessler, Heba Ismail

## Abstract

Understanding how connective tissue cells respond to mechanical stimulation is important to human health and disease processes in musculoskeletal diseases. Injury to articular cartilage is a key risk factor in predisposition to osteoarthritis. Here we identified a ubiquitin signature that is unique to injured articular cartilage tissue (the “injury ubiquitinome”). A total of 408 ubiquitinated peptides mapped to 114 proteins were identified, with an enrichment of ubiquitinated peptides of proteins involved in protein processing in the endoplasmic reticulum(ER), also known as the ER-associated degradation(ERAD) response, including YOD1, BRCC3, ATXN3 and USP5 as well as the ER stress regulators, RAD23B, VCP/p97 and Ubiquilin 1. Enrichment of these proteins suggested an injury-induced ER stress response and, for instance, ER stress markers DDIT3/CHOP and BIP/GRP78 were upregulated following cartilage injury on the protein and gene expression levels. Similar ER stress induction was also observed in response to tail fin injury in zebrafish larvae, suggesting a generic response to tissue injury. Furthermore, a rapid increase in global DUB activity following injury and significant activity in human osteoarthritic cartilage was observed using DUB specific activity probes. Inhibition of DUBs using a broad-spectrum inhibitor caused a reduction in the injury-induced inflammatory response in a zebrafish tail fin injury model. These results implicate the involvement of ubiquitination events and activation of a set of DUBs and ER stress regulators in cellular responses to cartilage tissue injury and osteoarthritis. This link through the ERAD pathway makes this protein set attractive for further investigation in *in vivo* models of tissue injury and for targeting in osteoarthritis and related musculoskeletal diseases.

## 1. Introduction

Articulated joints guarantee the smooth movement of mammals. Articular cartilage serves as a cushion protecting the surfaces of bones in the joints by preventing friction and attrition. Mechanical injury to articular cartilage results in chronic inflammation, pain and predisposes to tissue damage and loss of joint function. Progressive deterioration of the cartilage is the hallmark feature of osteoarthritis. Although multiple efforts have been made to understand how injury leads to osteoarthritis [1], it is still unknown how cells sense injury and what are the key cellular mechanism driving the damage.

Our previous work has shown that mechanical injury to articular cartilage activates within seconds the intracellular signalling pathways characteristic of the inflammatory response including TAK1[2], NFkB, MAP kinases [2–5], and Src, followed by induction of genes characteristic of inflammation [5]. Cartilage is an avascular tissue, so connective tissue cells alone must directly sense the injury and respond with inflammatory signalling and gene induction. Interestingly, the response is not restricted to the injury site and propagates through the tissue with time [2]. To identify the upstream mechanism, we have recently discovered that injury caused an overall increase in Lysine 63 polyubiquitination in porcine and murine cartilage [2].

Protein ubiquitination is an upstream mechanism of post-translational modifications which is emerging as a critical regulator of inflammatory signalling pathways [6]. Ubiquitination is a reversible and dynamic reaction tightly controlled by the opposing actions of ubiquitin ligases and deubiquitinating enzymes (DUBs) [6]. DUBs are cysteine and metallo-proteases that work by reversing and editing ubiquitination in a specific manner and play several roles in ubiquitin homeostasis and negative regulation of ubiquitin signalling [Reviewed in 7]. DUB activity can be modulated by further posttranslational modifications including phosphorylation and ubiquitination. The presence of at least one ubiquitin binding sites in DUBs facilitates their ubiquitination particularly when in complex with E3 ligases [Reviewed in 7]. Ubiquitination of DUBs could regulate their catalytic activity by either competing with binding of ubiquitinated substrates especially when associated with an E3 ligase, or by enhancing activity through activating conformational changes [8–10].

At the cellular function level, DUBs contribute to a plethora of cellular functions including cell signalling and modulation of ER stress. DUBs deregulation contributes to various disorders and age-related tissue degeneration [11–13]. A number of DUBs are reported to control stress including YOD1 and ATXN3. YOD1 and ATXN3 are highly conserved DUBs that are involved in the regulation of ERAD pathway by facilitating protein quality control through the ATPase VCP [14]. YOD1 is recently reported to be involved in inflammation through interaction with the E3 ligase TRAF6 and regulation of the inflammatory cytokine IL1 through competing with the p62 NFkB axis[15]. YOD1 attenuates neurogenic proteotoxicity through its deubiquitinating activity[16]. The role of ATXN3 in neurodegeneration is also reported through regulation of ER stress [17, 18].

The endoplasmic reticulum (ER) is the site for biosynthesis for many cellular components. The ER controls crucial cellular functions, including protein synthesis, lipid synthesis, Ca^++^ storage and release as well as Golgi and mitochondrial biogenesis. One of its key functions is protein quality control. Stress conditions that cause the protein import to exceed the protein folding capacity of ER trigger the unfolded protein response or UPR [19]. Failure of the ER to handle this stress response leads to cellular dysfunction, cell death, apoptosis and diseases including neurodegeneration, diabetes mellitus, cancer, etc [Reviewed in 20, 21, 22].

The UPR response is intended to restore the ER homeostasis through attenuation of protein translation and upregulation of ER-chaperone proteins CHOP, GRP78, and ATF3. Apoptosis is initiated when ER stress is not resolved [23, 24]. GRP78 plays a key role in detection and transduction of ER UPR signalling to downstream transcription factors. GRP78 also known as BiP/HSPA5 is normally bound to the ER stress sensors exist in the transmembrane; IRE1, ATF6 and PERK. When misfolded proteins accumulate in ER lumen, GRP78 dissociates from these sensors initiating downstream signalling through these three arms[24, 25].

In cartilage, the UPR response plays a physiological role in both chondrogenesis and chondrocytes maturation during endochondral ossification [26]. A prolonged ER stress response has also been reported to initiate chondrocyte death as identified in early onset osteoarthritis that is associated with several chondrodysplasias and collagenopathies characterised by prolonged UPR, ER stress and ERAD. An extensive and a comprehensive review on the role of ER stress and UPR in cartilage and OA is described by Hughes et al., 2017[26]. Understanding how UPR signalling is regulated in cartilage injury and related cartilage pathophysiology is thus key to therapeutic targets discovery in osteoarthritis and related musculoskeletal diseases.

Here we identify the unique ubiquitin signature of articular cartilage upon mechanical injury. We have observed differential ubiquitination patterns of YOD1 and ATXN3 deubiquitinases that modulated their DUB activity in injured and osteoarthritic articular cartilage. We also report a significant modulation of the ubiquitination pattern of the key ER stress regulators; VCP, RAD23B, and Ubiquilin 1 as well as the two E2 enzymes UBE2N and UBE2L3. Inhibition of DUBs exerted a significant delay in injury-induced inflammatory response in a zebrafish tail fin injury model as seen by reduced NFkB activation and reduced number of neutrophils recruited to the injury site in the presence of a broad spectrum DUB inhibitor. Our results highlight the importance of injury-induced ubiquitination events and DUB activation in regulating cellular responses that drive ER stress response. Our findings indicate that DUB activation contributes to the injury-induced inflammatory response and tissue damage, highlighting a possible mechanism for targeting in osteoarthritis.

## 2. Results

### 2.1. Mechanical injury to articular cartilage significantly modulates the cartilage ubiquitinome and proteome

We previously showed that injury to murine and porcine articular cartilage causes a significant increase in K63-polyubiquitination [2]. To identify the protein substrates that are differentially ubiquitinated in response to tissue injury tissue (the “injury ubiquitinome”), we used ubiquitin remnant motif analysis [27] of uninjured versus injured porcine articular cartilage. Based on our previous work, identified that the changes in ubiquitination post-injury are rapid and transient, with an optimum increase in ubiquitination at 30 minutes [2]. To identify the rapidly and differentially ubiquitinated peptides in response to articular cartilage injury, porcine cartilage was injured and biological triplicate samples were collected at 0, 10, and 30 minutes following injury. Injured tissues were lysed in a urea-containing buffer and subjected to tryptic digestion. Polyubiquitinated proteins were enriched from digested tissue lysates using anti-K-ε-GG antibody as described in methods. Samples were then analysed by LC–tandem MS (MS/MS) for quantitative profiles of non-redundant enrichment of ubiquitinated sequences/peptides as well as global proteome as previously described [28].

463 ubiquitinated peptides were detected. Data were filtered based on valid values and imputed missing values from normal distribution resulting in 408 ubiquitinated peptides (**Figure 1-Source Data 1**). Principal component analysis of control and injured time sets showed a good separation between samples based on injury stimulation (**Figure 1A**). Out of the 408 enriched ubiquitinated peptides, 201 with 3 charges and 189 with two charges, and 18 peptides with 4 charges were identified. Two different cut-off parameters were used to represent significantly enriched peptides post injury. Class A for FDR 0.05 and S0 0.5 and Class B for FDR 0.05 and S0 0.1. Volcano plots in **Figure 1B** show that a total of 170 ubiquitinated peptides were differentially enriched following 30 min post injury (Class A) and 200 enriched peptides (Class B) compared to 39 ubiquitinated peptides were enriched in the 10 min injury data set (Class A) while 52 ubiquitinated peptides were enriched in Class B (Figure 1-Source Data 1). The heat map in **Figure 1C** shows a group of significantly enriched ubiquitinated peptides that are common to both the 10 min and 30 min injury data sets, ranked by log2 fold of change in the 30 min-injury set. This group included 38 peptides mapped to 27 proteins for example RPS3, VCP, HSPA8 and ARL8.

**Figure 1:**
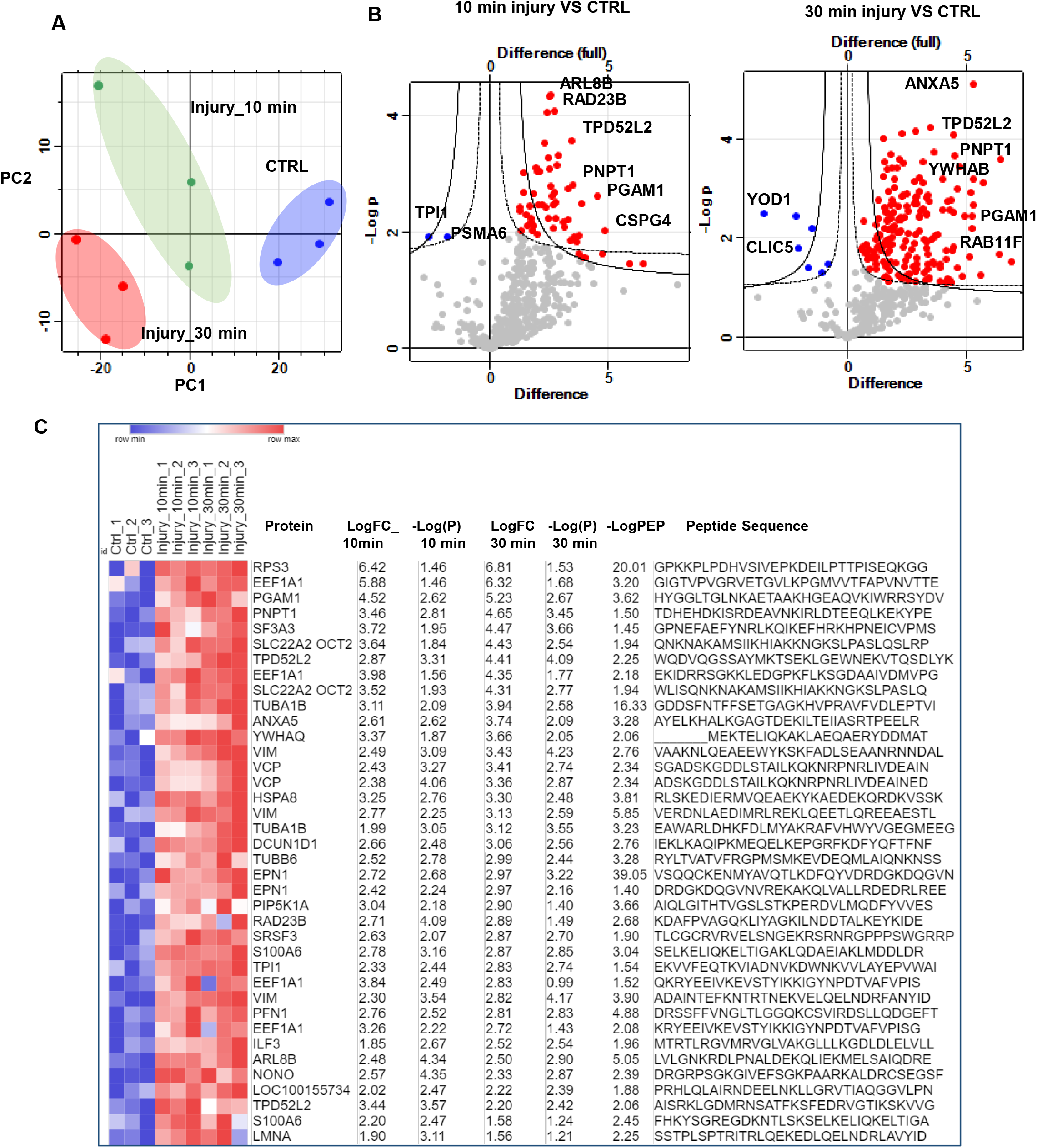
Ubiquitomics landscape of articular cartilage before and after mechanical injury. **A)** Principal component analysis(PCA) plot of ubiquitinated peptides intensity in control uninjured (CTRL) and injured articular cartilage samples 10 and 30 minutes post injury reveals strong clustering of samples by injury response in the three biological replicates. **B)** Volcano plot representation of ubiquitomics analysis of articular cartilage samples before (CTRL) and after injury (10 min and 30 min). **C)** Heat map of differentially ubiquitinated peptides in response to injury that are commonly detected after 10 and 30 minutes compared to uninjured control ranked by log2 fold of change(LogFC). Statistics by unpaired t-test.

These results highlight that injury to articular cartilage causes a rapid change in protein ubiquitination patterns of a specific set of cellular proteins. This change occurs very rapidly within 10 min and is optimum at 30 min.

The enriched ubiquitinated peptides post injury were mapped to 115 proteins. Enriched pathways were detected using STRING v11.0 database [29]. Examples of the top enriched pathways are shown in **Figure 2A** bar chart. The top enriched KEGG pathways included Tight junction (9 proteins), ‘’Protein processing in endoplasmic reticulum’’ (7 proteins), Endocytosis (6 proteins), carbon metabolism (5 proteins), biosynthesis of amino acids (5 proteins), glycolysis / gluconeogenesis (4 proteins), Ribosome (5 proteins) and carbon metabolism (5 proteins). Reactome pathways analysis showed a significant enrichment of vesicle mediated transport (20 proteins), membrane trafficking, clathrin mediated endocytosis, translocation of GLUT4 to membrane and Josephin domain DUBs (5 proteins), and metabolism of proteins. Details of the enriched pathways and linked proteins are shown in **(Figure 2-Source Data 1)**. Mapped proteins were further analysed for protein-protein interactions based on experimental evidence using STRING database. Disconnected nodes were excluded and only directly interacting proteins based on experimental evidence are shown. Out of 115 mapped proteins, 68 proteins showed node connections based on experimental evidence with PPI enrichment p-value= 2.2e-05. Those were separated into 5 main clusters based on protein function including ERAD, carbon metabolism and ribosome protein clusters (**Figure 2B**).

**Figure 2:**
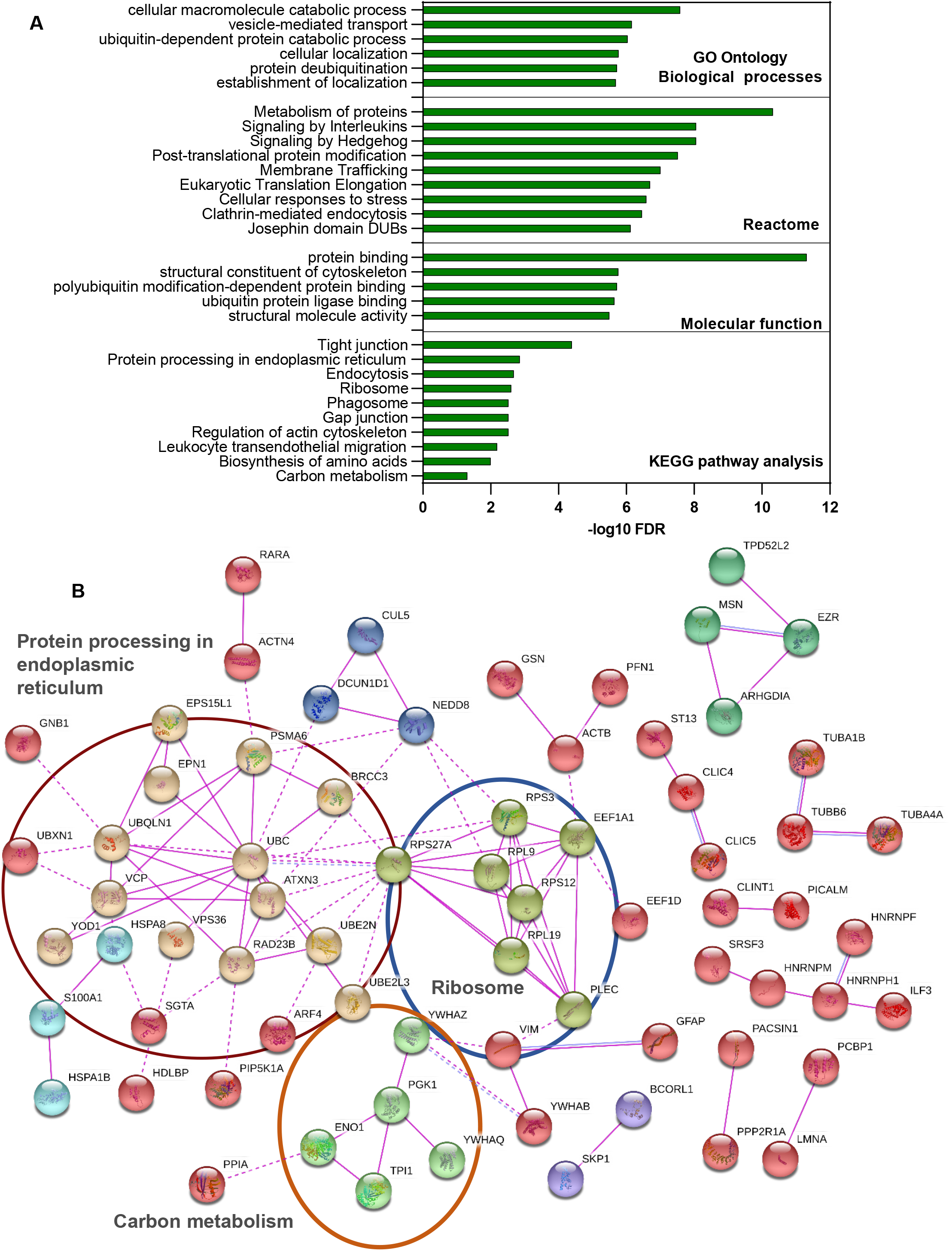
Protein network and enriched pathway analysis of differentially ubiquitinated proteins in articular cartilage post injury. **(A)** Pathway analysis of enriched ubiquitinated proteins modulated in response to injury using KEEG, Reactome and Go ontology pathway analysis databases. **B)** ubiquitinated peptides identified in cartilage post injury were mapped to proteins and were further analyzed for direct protein-protein interactions using STRING database. Disconnected nodes are hidden and only connections confirmed in experiments are shown.

Global proteome of cartilage lysates before and after injury was also analysed using mass spectrometric analysis as described in methods. 2450 proteins were detected across all samples (**Figure 3-Source Data 1)**). Data were filtered based on valid values and imputed missing values from normal distribution, resulting in 2394 annotated proteins. Class A significance (S0 0.5 & FDR0.01) of proteome showed enrichment of 564 proteins in cartilage at 30 min post injury compared to 63 proteins (Class A) were enriched in the10 min proteome data set. Class B significance (S0 0.5 & FDR 0.05) of proteome showed 1145 proteins enriched at 30 min while 201 proteins were enriched 10 min post injury (**Volcano plots in Figure 3A&B**). Heat map in **Figure 3C** shows the top 20 proteins that are commonly modulated post injury after 10 and 30 minutes. Enriched proteins (Class A) after 30 min injury were interrogated using the STRING database to identify protein-protein interaction and enriched pathways. The bar chart in Figure 3D show examples of enriched KEGG and Reactome pathways, including extracellular matrix organisation, protein metabolism and protein processing in endoplasmic reticulum (UGGT1, UBXN6, HSPA6, UBE2G2, SEC24C, RAD23B, ERO1L, DNAJB11, BAG1, PRKCSH and RAD23A).

**Figure 3:**
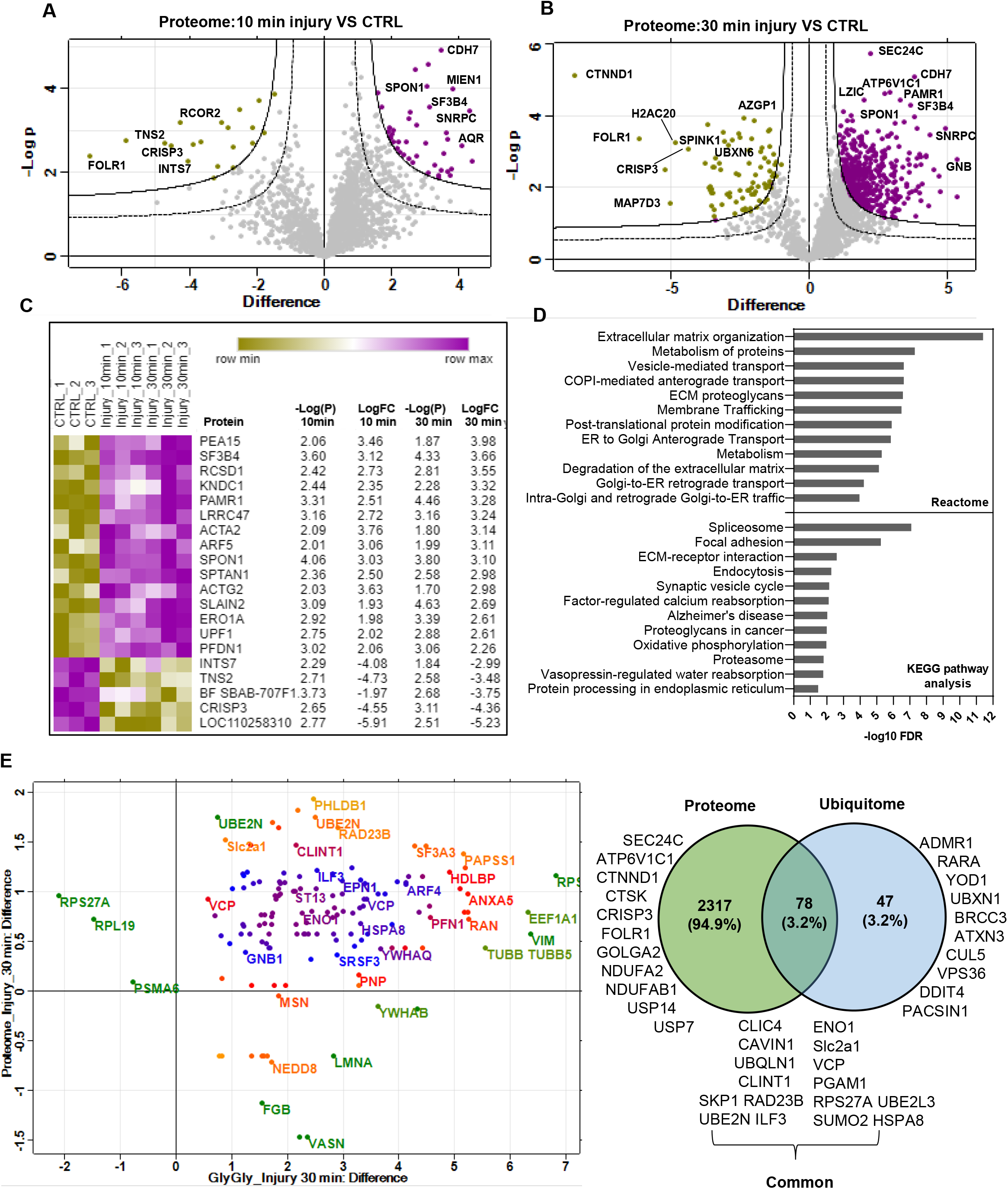
Proteome analysis of porcine articular cartilage tissue samples and comparison with ubiquitinome analysis. **A&B)** Volcano plots representation of proteomics analysis of articular cartilage samples before (CTRL) and after injury (10 min and 30 min). **C)** Heat map of top 20 proteins differentially regulated post injury that are commonly detected after 10 and 30 minutes are shown ranked by log fold of change in the three biological replicates. Statistics by unpaired t-test. **D)** Pathway analysis of enriched proteins in proteome analysis post injury using KEEG and Reactome STRING analysis databases. **E)** Cross analysis of cartilage ubiquitinome and proteome. Scatter plot shows proteins that are commonly detected in ubiquitinome and proteome 30 minutes post injury. Venn diagram shows a comparison of significantly enriched ubiquitinated proteins and total proteome 30 min post injury.

Cross analysis of the injury proteome and injury ubiquitinome showed that, of the 115 proteins that are significantly ubiquitinated post injury, 78 proteins were also enriched post injury in the proteome dataset (3.2% of total proteome) **(Figure 3E & Figure 3-Source Data 2))** while 34 proteins were uniquely enriched in ubiquitinome dataset post injury including YOD1, ADMR1, UBXN1, ATXN3 and DDIT4 proteins.

### 2.2. Cartilage injury induces ubiquitination of a protein subset of ER regulators and deubiquitinases involved in the regulation of the ERAD response

We interrogated the set of differentially ubiquitinated peptides post injury using STRING and identified a cluster of enriched peptides that are highly connected in a protein network. The proteins identified in this cluster were mapped to the GO term “response to cellular stress” and the KEGG pathway “protein processing in the endoplasmic reticulum” also known as the ERAD response. This cluster was also mapped to the GO terms ‘’ubiquitin protein ligase binding’’ and ‘’protein deubiquitination’’ (**Figure 2**). This protein cluster comprised the three deubiquitinases YOD1, BRCC3 and ATXN3, the two E2 ubiquitin ligases UBE2N and UBE2L3 as well as the ER stress regulators VCP/p97, RAD23B, Ubiquilin 1, UBXN1, SAR1A and HSPA8 proteins **(Figure 2B)**. Hereafter this set will be referred to as the ER/ DUB protein set. The identified ER/DUB set of proteins is well known for their involvement in the ERAD response [30–32], making this protein set attractive for further analysis.

The heat map in **Figure 4A** shows the ubiquitination pattern of this protein set after injury in all replicates compared to uninjured controls. We have validated and confirmed the differential ubiquitination of some proteins in this ER/DUBs protein set in response to injury using ubiquitin enrichment pulldown assays followed by western blotting as described in methods and in **Figure 4B**. The 3 DUBs BRCC3, ATXN3 and USP5 ubiquitination is enhanced post injury (**Figure 4C**). YOD1 is enriched in ubiquitinated fraction in uninjured controls and was significantly deubiquitinated post injury (**Figure 4C; right panel**). **Figure 4E** shows the enrichment of the three ER regulators VCP, ubiquilin 1 and RAD23B in ubiquitinated proteins pull down 30 min post injury compared to the uninjured control. We validated the ubiquitination of the two E2 ubiquitin conjugating enzymes UBE2L3 and UBE2N as shown in **Figure 4E**. Figure 4F shows the protein levels of this ER/DUB set in input tissue lysates before and after injury used for ubiquitin enrichment assay compared to b-actin as a loading control in **Figure 4A-E**.

**Figure 4:**
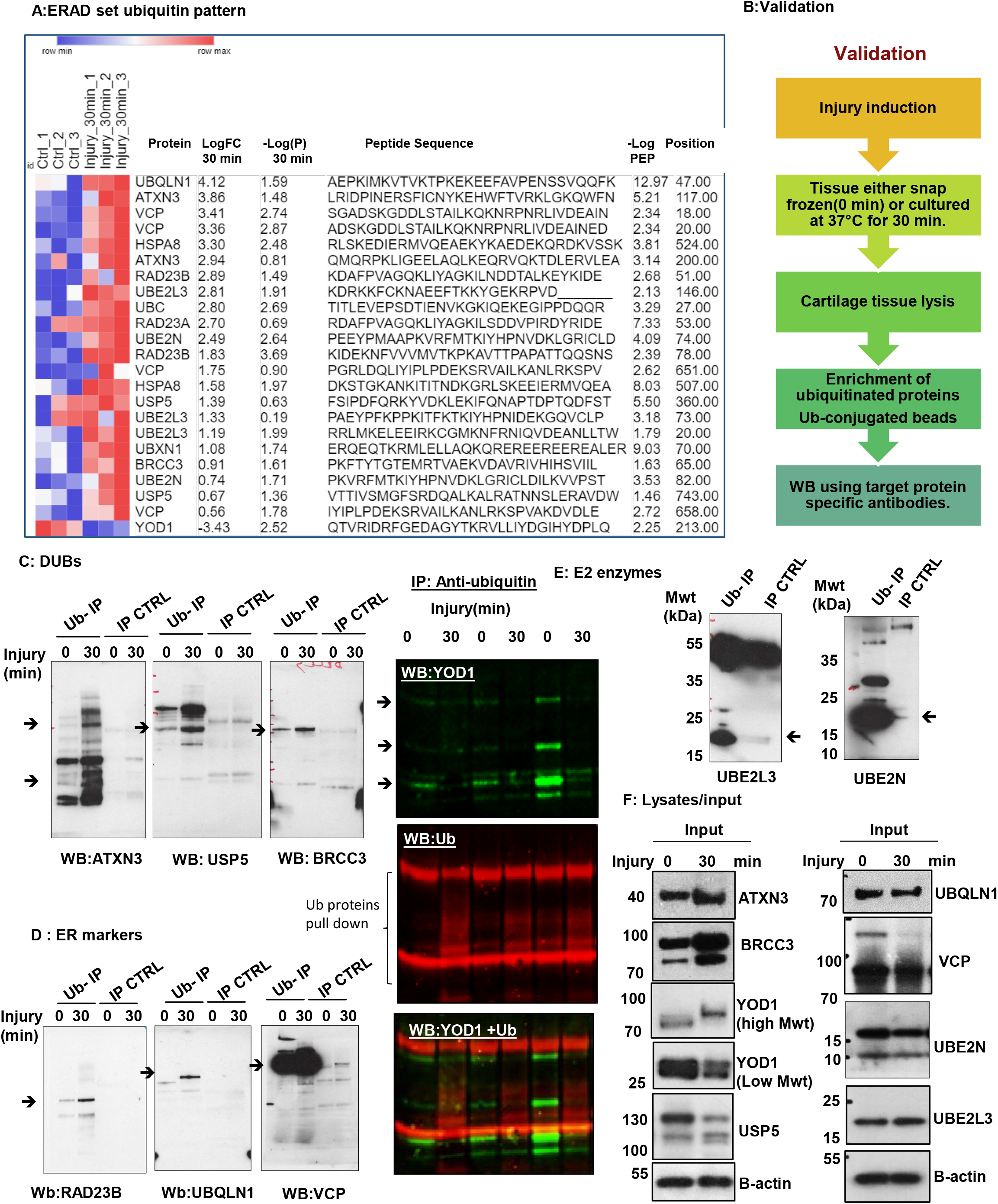
Key regulators of ERAD pathway are differentially ubiquitinated in response to articular cartilage injury. **A)** Heat map of differentially ubiquitinated peptides of ERAD set proteins in response to injury (30 minutes) compared to uninjured control ranked by log2 fold of change in three biological replicates. Statistics by unpaired t-test. **B)** A summary of the validation approach used to confirm ubiquitomics analysis. **(C-E)** Porcine cartilage was explanted to induce injury and either snap frozen(0 min) or cultured at 37°C for 30 min. Cartilage was then lysed as described in methods. Ubiquitinated proteins were enriched using anti-ubiquitin conjugated agarose beads(Ub-IP) and agarose beads were used as a control(IP CTRL). After immunoprecipitation, Immunocomplexes were analysed by western blotting using target protein specific antibodies for DUBs proteins in **(C)** ATXN3, USP5, BRCC3 and YOD1, **(D)** ER stress markers (VCP, RAD23B, UBQLN1), or **(E)** E2 enzymes UBE2L3 or UBE2N. Arrows indicate the enriched specific protein. Shown are blots representative of three biological replicates. **(F)** Protein level of candidate proteins were analysed in input tissue lysates samples. B-actin was used as internal loading control.

These results validate the ubiquitomics mass spectrometry analysis and confirms the differential ubiquitination of the ER/DUB protein set post injury. Data shown here suggests a significant involvement of ER stress response and DUB related proteins in cellular responses to mechanical injury to articular cartilage.

### 2.3. Injury to articular cartilage and zebrafish embryos causes induction of an ER stress response

Having observed that articular cartilage injury induces ubiquitination of the key ER stress regulators VCP, Ubiquilin 1 and RAD23B (**Figure 4E**), we hypothesized this may reflect the induction of ER stress response upon injury. To test the direct link between cartilage injury and ER stress, we interrogated our previously published microarray data of murine hip cartilage before and after 4hrs of tissue injury (**GEO accession number: GSE155892**) for gene expression levels of ER stress genes. We observed a significant upregulation in expression levels of ER stress related mouse genes including Atf3, Atf4, Atf5, Ddit4, Trib3, Hspa1a &b and Ddit3/CHOP. A marginal increase was observed in the levels of Xbp1, HSPA8 and Ern1 genes post injury **(Figure 5A)**. We then investigated the gene and protein expression of ER stress markers GRP78, XBP1 and CHOP following injury to porcine articular cartilage. Porcine cartilage tissue was injured as for the indicated times then processed either for RNA or protein extraction as described in methods. Gene expression levels of CHOP, GRP78 and XBP1 ER stress genes were detected using real time RT-qPCR. GAPDH was used as housekeeping gene. **Figure 5B** shows a transient elevation of ER stress genes expression following mechanical injury with a peak of induction between 2-4hrs. This increase reset to basal levels by overnight incubation of cartilage following injury. **Figure 5C** shows differential protein levels of CHOP 24hrs following cartilage injury. Interestingly, we have noticed that injury lowered expression levels of CHOP lower molecular weight form (27 kDa) and induced a higher molecular weight form after 24hrs of injury. This may be a super shift form of CHOP protein caused by its ubiquitination and indicating its activity. Ubiquitination of CHOP protein causes its shift to 50 kDa was previously reported [33]. Induction of Xbp-1 gene expression as well as its splicing is also a marker of ER stress activation [34]. We have noticed that Xbp-1 spliced variant is induced following tissue injury in articular cartilage as early as 30 minutes (**Figure 5D**).

**Figure 5:**
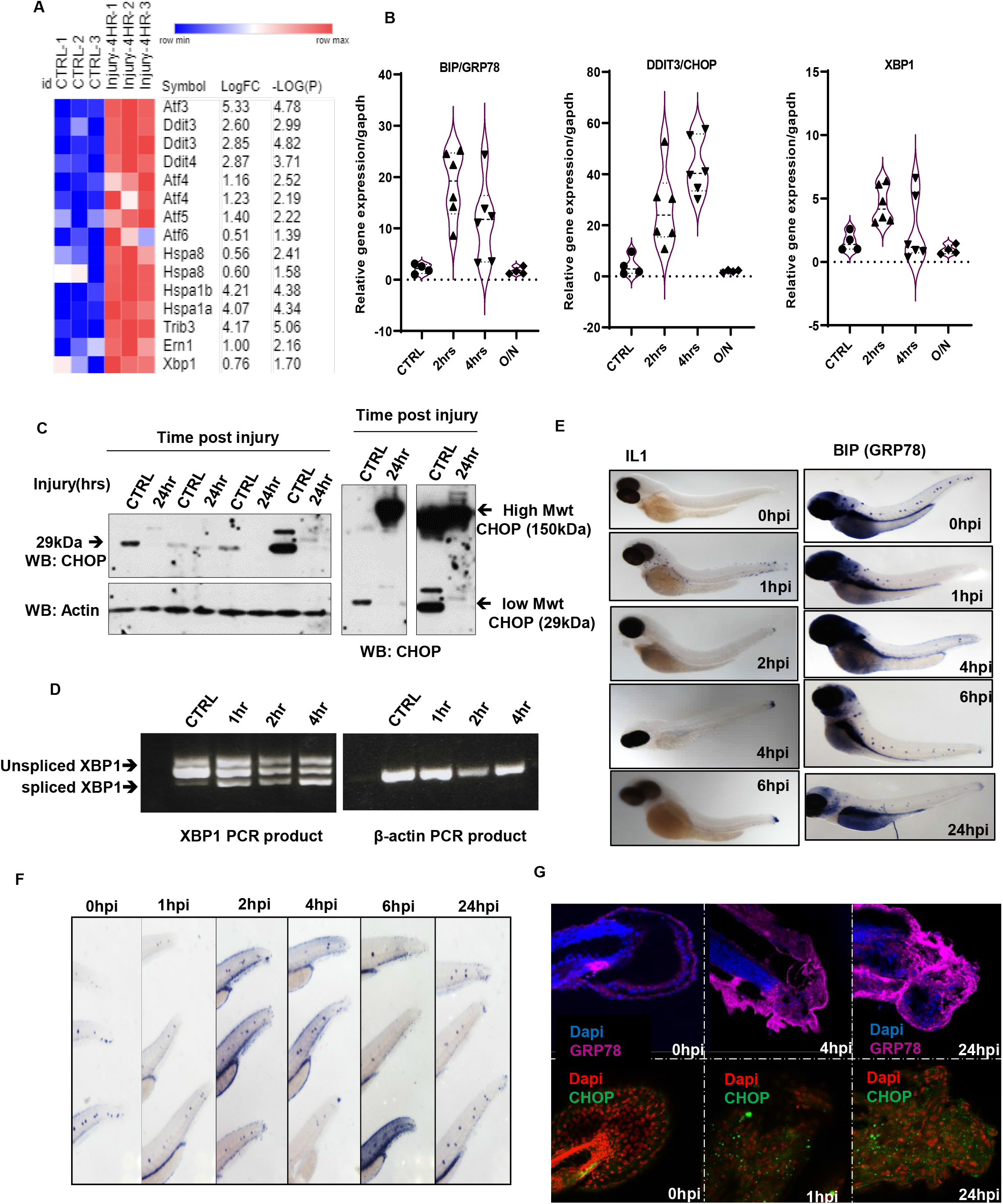
Injury upregulates the ER stress markers in murine and porcine articular cartilage as well as in zebrafish embryos. **(A)** Heat map of ER stress genes in murine hip cartilage before and after injury.ER stress genes expression levels were retrieved from our previously published murine hip injury microarray data set (GEO: GSE155892). Statistics by unpaired t-test. **(B)** RT-qPCR of BIP, DDIT3 and XBP1 genes in porcine articular cartilage post injury. **(C)** CHOP protein levels in porcine cartilage lysates before and after 24hrs post injury(4 biological replicates). **(D)** PCR products of XBP1 from porcine cartilage before and after injury were analyzed by agarose gel electrophoresis. **(E).** In situ hybridization of IL1(positive control) and GRP78 RNA probes in 3dpf zebrafish embryos before and after tail fin injury. Images were obtained using Stereomicroscope-integrated colored camera. **(F)** GRP78 RNA staining in zebrafish tails before and after injury (purple stain) **(G)** immunofluorescence analysis of GRP78 and CHOP proteins in zebrafish injury. DAPI for molecular stain. Images z stacks were obtained using A1 Nikon confocal microscope and are shown after maximum intensity projection.

Our previous work showed that injury responses are generic to connective tissues in porcine, murine and zebrafish tissues (**[2, 35, 36] & unpublished data**). To test if injury caused an induction of ER stress response *in vivo*, we have used a well-established tail fin injury model of developing zebrafish embryos. Injury was induced by transection of zebrafish embryo tails at 3days post fertilisation as described in methods. We evaluated the induction of ER stress by increased levels of GRP78 mRNA and GRP78 and CHOP proteins at the injury site using RNA *in situ* hybridisation and immunofluorescence respectively (**Figure 5E&F**). We have observed that GRP78 expression in uninjured control embryos is localised around neuromasts while expression levels are increased around the injury site as early as 2 hours post injury. IL1 RNA probe was used as a positive control for injury responses in zebrafish embryos (**Figure 5E left panel**). We also observed increased protein levels of CHOP and GRP78 at the tail fin injury site using immunofluorescence **(Figure 5G).**

Taken together, these data indicate that mechanical injury to articular cartilage directly causes an induced ER stress response reflected by increased gene and protein expression levels of ER stress markers. The data also indicates that the responses to tissue injury are generic and the effect on ER stress is also observed *in vivo* as well as ex-vivo models of tissue injury.

To investigate if the expression pattern of ER stress responsive genes is modulated in osteoarthritis, we interrogated the publicly available Gene Expression Omnibus (GEO) RNA sequencing data available for osteoarthritic tissue samples (synovium and cartilage) (**Supplementary Figure** 1). Interestingly we have observed a significant reduction in the expression of ER transcription factors CHOP, ATF3, XBP1 while ATF6 and ATF4 transcription factors levels were not changed. ER stress genes HSPA5, ERN1, were also significantly reduced in OA cartilage compared to normal. DERL1,2, and 3 genes were not changed in diseased samples. Interestingly, HSPA8 levels were significantly increased in OA cartilage relative to normal samples. HSPA8 was reported before to be downregulated in response to ER stress (REF). To test if the profile of the key transcription factors ATF3, CHOP and XBP1 is affected in OA synovium similar to what we observed in cartilage, we analysed 3 RNA sequencing studies (GDS2126, GDS5401 and GDS5403) comparing OA synovial biopsies with normal synovial biopsies. Interestingly the expression levels of the three key genes are markedly reduced in OA samples compared to normal counterparts consistent with the same observation in cartilage (S**upplementary Figure 1B**).

These data highlight a significant disruption of expression of genes necessary for ER homoeostasis which may indicate that ER homeostasis is affected in osteoarthritis.

### 2.4. A rapid increase in global DUB activity following injury to porcine cartilage tissue and detection of significant DUB activity in human osteoarthritic cartilage

To investigate if DUBs are active in injured or diseased cartilage, we have analysed the global DUB activity in porcine and human articular cartilage. Using DUB specific activity probes, we compared the DUB activity in injured and control porcine articular cartilage samples. Interestingly, we observed an increase in DUB activity following injury as shown in **Figure 6A.** At the same time, other DUBs appear to be enriched in the control set compared to the injury set, indicating possible deactivation of another group of DUBs upon injury.

**Figure 6:**
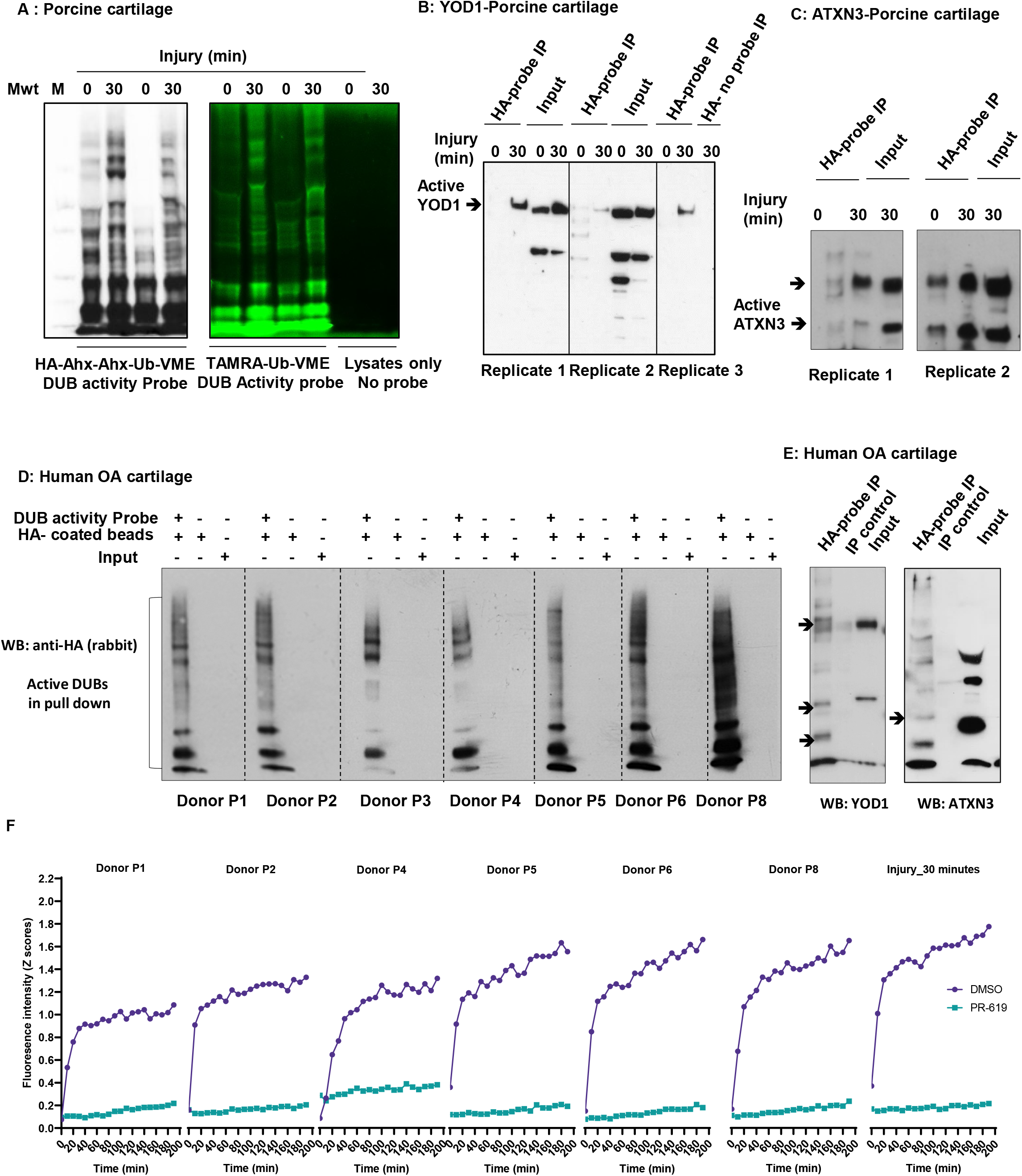
DUBs are active in injured and osteoarthritic cartilage. **(A)** Porcine articular cartilage lysates (control and injured) were incubated with the DUB activity probes HA-Ahx-Ahx-Ub-VME or TAMRA-Ub-VME for 30 min at 37°C. Western blotting of HA-Ahx-Ahx-Ub-VME DUB activity reactions of cartilage lysates using anti-HA antibody (Left panel) and In-gel fluorescence scan of TAMRA-Ub-VME deubiquitination reactions was obtained with a XX6 gel imaging system (Syngene) (right panel)(resolution= 100 μm and exposure time of 60s, λex/λem= 550/590 nm). Porcine cartilage **(B-C)** and human OA cartilage **(D-E)** obtained and lysed as described in methods. 50 μg of lysates were reacted with the DUB activity probe HA-Ahx-Ahx-Ub-VME for 45 minutes at 37°C then subjected to immunoprecipitation of active DUBs using mouse anti-HA antibody-conjugated magnetic beads. Immuno-complexes were analysed by western blotting and compared with input and IP control using DUB specific antibodies YOD1 **(B),** ATXN3**(C),** and YOD1 and ATXN3 for OA cartilage **(E). D)** active DUBs pulldown in OA cartilage tissue lysates from 7 different donors. Were analysed by western blotting using rabbit anti-HA antibody. **F)** UBRH110 DUB activity assay of human OA cartilage and porcine injured cartilage in the presence or absence of PR-619 DUB inhibitor

As described above, we have observed a differential ubiquitination pattern of YOD1 (Lysine 213) and ATXN3 (Lysine 117 and Lysine 200) in response to cartilage injury (**Figure 4C**). Ubiquitination of ATXN3 at Lysine 200 and YOD1 at Lysine 213/257 have not been reported before. It is well described that deubiquitinase activity can itself be modulated by ubiquitination [8–10]. This implicates a possible modulation of these DUB activities in our experiments following cartilage injury. To analyse if injury-induced ubiquitination of these identified DUBs in response to injury modulates their protease activity, we used DUB activity specific probed reactions, with active DUBs pulled-down and analysed by western blotting using DUB specific antibodies. Briefly, porcine cartilage tissue lysates (before and after injury) were reacted with HA-tagged DUB activity probe (HA-Ahx-Ahx-Ub-VME) for 45 min at 37°C. Reactions were then stopped and active DUBs that bind to the activity probe were pulled down using HA antibodies. Pull down complexes were then analysed by western blotting using anti-YOD1 and anti-ATXN3 specific antibodies. We observed that YOD1 and ATXN3 were pulled down in active DUBs fractions following injury (**Figure 6B &5C)** implicating that injury modulated-ubiquitination of these DUBs enhances their protease activity. We also observed USP5 activation post injury however, USP5 blot showed a nonspecific band above the predicted size of UPS5 but also showed enrichment of active USP5 in the pull down compared to IP control (Figure 6-Figure **Supplement 1**).

To confirm disease relevance of these findings, we investigated the DUB activity in tissue lysates of human OA cartilage using DUB activity assays (as described above). Osteoarthritic cartilage lysates from 7 human donors were reacted with DUB activity probes as described in methods and samples were treated as described for porcine cartilage. Western blotting of HA pulldown showed an enrichment of active DUBs in tissue lysates-activity probe immunoprecipitated (probe-IP) samples compared to beads control (CTRL-IP) indicating a significant level of DUB activity in those samples **(Figure 6D)**. YOD1 and ATXN3 were detected in active DUBs pull down fraction compared to control (**Figure 6E**) indicating that they are activated in osteoarthritic cartilage. To quantify the DUBs activity in tissue lysates we also used a quenched ubiquitin Rhodamine 110 substrate. We have detected the DUB activity in tissue lysates after injury and in osteoarthritic cartilage of 6 donors. The substrate fluorescence intensity increased over time reflecting enhanced DUBs activity **(Figure 6F)**. Pre-incubation of tissue lysates with PR-619 inhibitor abolished this activity confirming the activity we detected is related to DUBs in the lysates. The activity of DUBs in human OA and porcine injured cartilage was significantly suppressed when lysates are incubated with the broad-spectrum DUB inhibitor PR-619 confirming the above results. These findings demonstrate a global activation of DUBs in osteoarthritic and injured cartilage and enrichment of active ATXN3 and YOD1.

### 2.5. DUBs inhibition significantly reduces/delays inflammatory response to localised tissue injury in zebrafish embryos

We previously reported that mechanical injury to connective tissue as cartilage and synovium induces inflammatory signalling pathways, including MAPK and NFkB as well as induction of injury responsive genes [2, 5]. To test the role of injury-activated DUBs in *in vivo* injury-induced inflammatory response, we have used a well-established model of tail fin injury in Zebrafish. Zebrafish transparent embryos provide an excellent tool for advanced imaging of fluorescent transgenic reporters to monitor the dynamics of injury responses *in vivo*.

Zebrafish tail fin injury causes an activation of the inflammatory response as seen by activation of inflammatory NFkB signalling pathway [37] and neutrophil recruitment to the injury site [38]. We used MPO transgenic line in which neutrophils are fluorescently labelled green and their migration post injury monitored by wide field fluorescence microscopy. DUB activity was modulated using the broad-spectrum inhibitor PR-619. Zebrafish embryos were pre-treated with PR-619 prior to injury. Injury was induced as described in the methods and in Figure 7A then neutrophils migration was monitored at injury site under fluorescent streomicroscope. As shown in **Figure 7 and Figure 7-Source Data 1&2**, pre-treatment of 3dpf embryos with 50μM of the broad spectrum DUB inhibitor PR-619 for 6hrs or 24hrs before injury caused a significant delay in inflammatory response as seen by the reduced number of neutrophils recruited to injury site after 2hrs post wounding compared to untreated controls (**Figure 7B &C**). PR-619 treatment also caused reduction in neutrophil migration using concentrations ranging from 10-5μM (Figure 7-**Supplement Figure 1**). Live imaging of neutrophils migration post injury and tracking analysis showed a significant effect of PR-619 on the velocity of neutrophil migration to the wounding site while no effect on displacement (**Figure 7D&E, Movies 1 & 2**).

**Figure 7:**
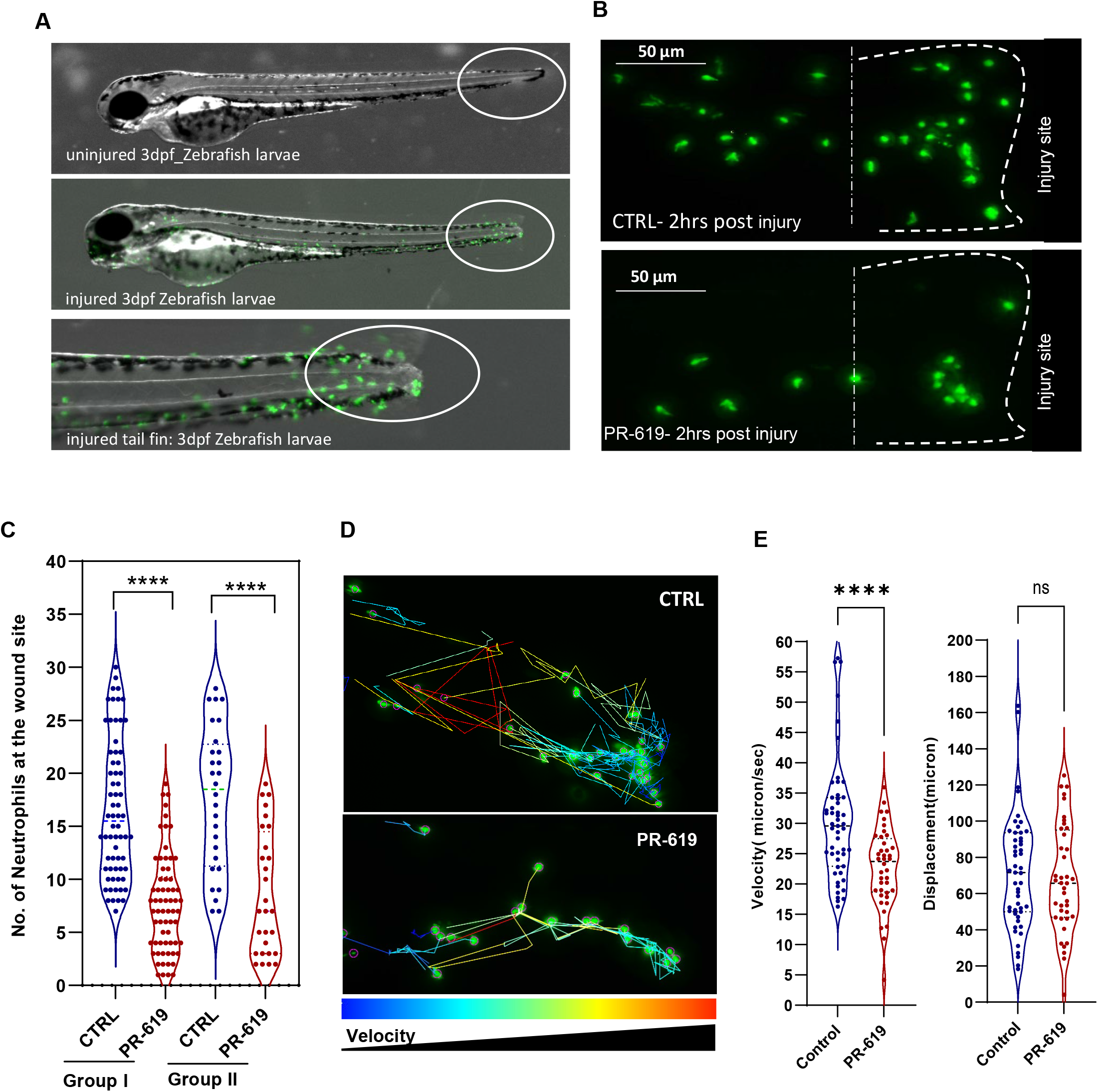
Effect of DUBs inhibition on injury induced-inflammatory response detected by neutrophil recruitment at wound site following tail fin injury in 3dpf Zebrafish Larvae. Tg(mpx:GFP)i114 neutrophil reporter line larvae(3dpf) were pretreated with either DMSO as control or 50 μM PR-619 inhibitor. Neutrophils (Green) were then monitored by live time lapse imaging using wide field microscope or counted at the wound site 2hrs following injury using fluorescent stereomicroscope. **(A)** representative stereomicroscopic images of uninjured and injured 3 days post fertilisation (3dpf) zebrafish with neutrophils at injury site. **(B)** Representative max intensity projection of live imaging stacks of neutrophils 2hrs post injury. **(C)** 3dpf larvae were treated for 6 hrs before tail fin injury as in Group I or 18 hrs as in Group II. Neutrophils were then counted at the wound site 2hrs following injury. Data shown are from 4 biological replicates(9-15 larvae/treatment). **D-E:** tracking of neutrophil migration was performed using FiJi plugin Trackmate. **(D)** Individual neutrophil tracks (from two individual larvae) in wounded 3dpf *Tg*(*mpx:GFP*) larvae treated with DMSO (CTRL) or PR-619. Color intensity reflects speed of neutrophils migration. **(E)** The effects of 50 μM PR-619 on neutrophil velocity and displacement when tracked in 3dpf *Tg*(*mpx:GFP*)*i114* wounded larvae, relative to DMSO controls [neutrophil tracks generated from 6 individual wounded larvae/treatment]. n.s., not significant; \*\*\*\**P*<0.0001.

We also used a recently established high-sensitivity bi-directional NFkB reporter transgenic zebrafish line [37] to monitor NFKB activation post injury using spinning disc confocal microscopy. DUB activity was modulated using the broad-spectrum inhibitor PR-619. Zebrafish embryos were pre-treated with PR-619 prior to injury. Injury was induced as described in the methods. NFkB activation was monitored by time-lapse confocal imaging of the injured tail fin area at intervals of 30 minutes for 18 hours. Images were analysed in ImageJ. We observed that DUBs inhibition with PR-619 caused a significant reduction of NFkB activation post injury at the wounding site (**Figure 8A, lower panel &Movie 4**) compared to untreated larvae (**Figure A, top panel &Movie 3**). **Figure C** shows NFkB activation over time as measured by mean fluorescence intensity of GFP reporter at wound site (**Left panel**) and bead area (**right panel**) in PR-619 treated and control untreated larvae. NFkB activation at notochord beads area was highly augmented and was also accompanied by reduced beads area in PR-619 treated fish compared with control (**Figure 8B &Movies 5 & 6**). Notochord bead areas are extrusions that are reported to be involved in repair of tail fin tissue post injury [39] highlighting a possible role of DUBs and NFkB activation in this tissue repair/regeneration mechanism.

**Figure 8:**
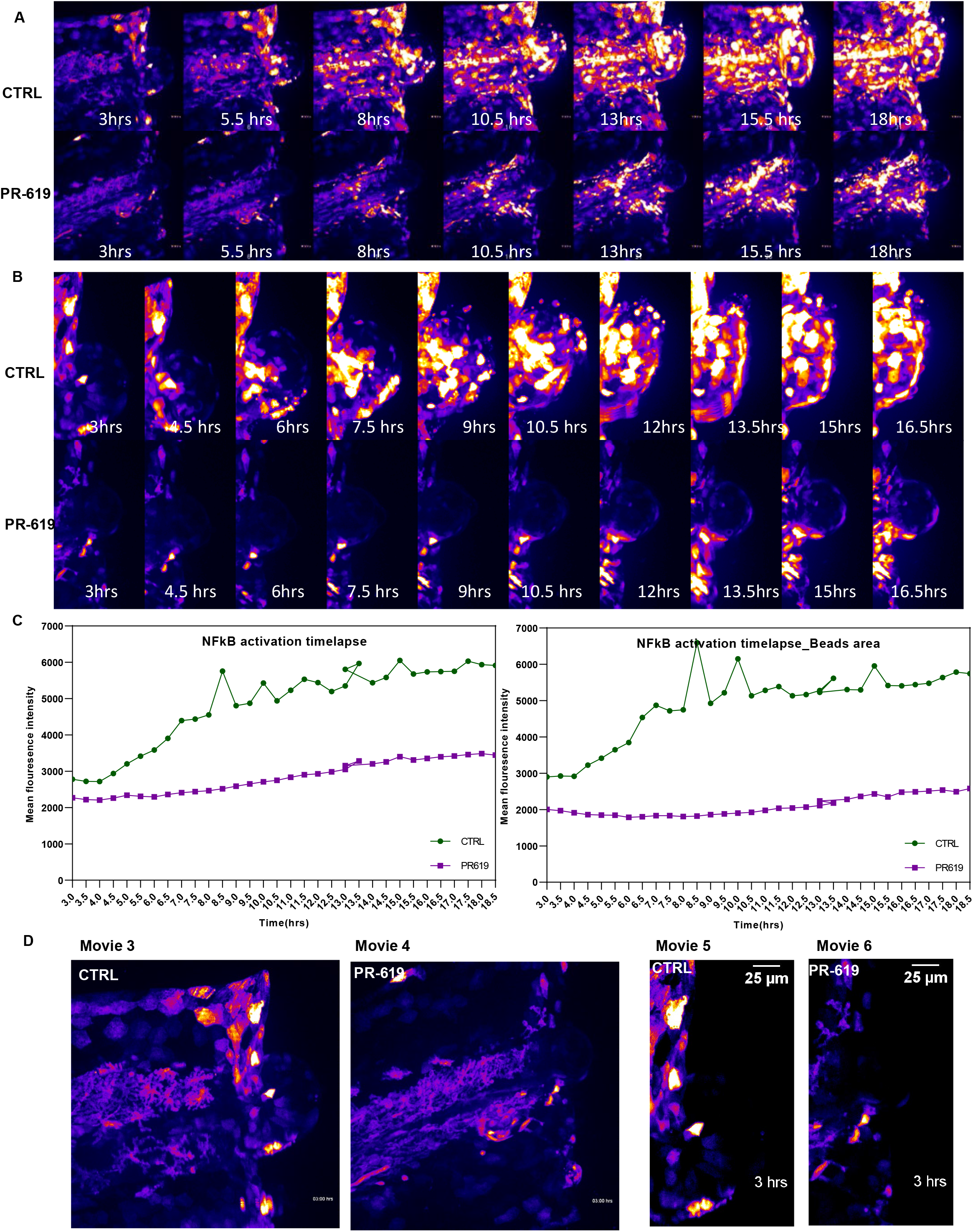
Effect of DUBs inhibition on injury-induced inflammatory response detected by NFkB activation at wound site following tail fin injury in 3dpf Zebrafish Larvae. Tg(8×Hs.NFκB:GFP,Luciferase) NFkB reporter line larvae(3dpf) were pre-treated with either DMSO as control(CTRL) or 50 μM PR-619 inhibitor for 6 hrs before tail fin injury. Injury was induced as described in methods and live images of wound sites was obtained using spinning disk confocal microscope every 30 min for 18 hrs (6 individual larvae/group)**. (A)** Time-lapse images derived from movies 1A and 1B show NF-κB activation after tailfin injury in control (top panel) compared to PR-619 treated larvae (lower panel). **(B)** Time lapse images of tail fin notochord bead area post injury in control and PR-619 treated fish. Images derived from movies 2A and 2B **(C)** Quantification of fluorescence intensity of NFkB activation post injury in control and treated tail fin and notochord bead areas.) in vivo visualization of NFkB activation in Tg(8xHs.NFκB:GFP,Luciferase) embryos post tail fin injury. GFP fluorescence is shown as false-coloured heat map. Fish anterior is to the left. Scale bar: 70μm

Taken together, we have demonstrated that inhibiting DUB activity using the broad-spectrum DUB inhibitor PR-619 reduces the inflammatory response following injury in a zebrafish model, as seen by reduced number of neutrophils recruited to wound site and reduced injury-induced NFkB activation. These findings implicate DUBs in modulation of inflammatory cellular responses to localised connective tissue injury.

## Discussion

Injury to articular cartilage is a key risk factor in the initiation and progression of osteoarthritis, a degenerative disease with no modifying therapies identified to date [Reviewed in 40, 41]. Identifying the early cellular responses to injury would facilitate understanding the early events that drive the tissue damage and consequently the loss of function in this disease. Our data highlights a significant ubiquitin signature of articular cartilage modulated by mechanical injury. Proteins significantly enriched were key regulators in ubiquitin system and ERAD response. However, whilst the ubiquitin system is implicated in several diseases, to date no studies have focused on its role in articular cartilage damage and in osteoarthritis. We here present for the first time the unique ubiquitin signature of cartilage injury and report the possible involvement of ER stress in modulating cellular responses that cause injury–induced tissue damage in articular cartilage.

In this study, we propose a model of early responses to tissue injury that causes enhanced ubiquitination events and involves key proteins that regulate ER stress and other cellular responses. The key players of ER stress VCP, Ubiquilin 1, RAD23B together with the ER associated DUBs YOD1 and ATXN3 were differentially ubiquitinated following injury in our ubiquitomics experiments **(Figures 1&4)**. These results were validated by ubiquitin enrichment pull down assays followed by western blotting. Interestingly, these ubiquitination events were very rapid and affected downstream proteins involved in ER stress response.

Through the ER stress adaptive response, the cell maintains physiological homeostasis and survival in response to stresses, including metabolic disturbance, reduced calcium levels, augmented oxidative stress and the accumulation of unfolded or misfolded proteins. This response causes a translational attenuation accompanied by activation of the ERAD response and activation of chaperones gene expression. However, excessive and prolonged activation of the ER stress response causes cellular death [42, 43]. Interrogating our previously published microarray data of murine hip cartilage, we observed that injury to hip cartilage causes a significant upregulation in expression levels of ER stress related mouse genes including Atf3, Atf4, Atf5, Ddit4, Trib3, Hspa1a &b and Ddit3/CHOP **(Figure 5A)**. We also observed that injury to porcine cartilage induces gene expression levels of ER stress genes GRP78/BIP, XBP1, and CHOP. CHOP and GRP78 proteins were upregulated after injury in zebrafish embryos as well as at gene expression level in porcine cartilage following injury (**Figure 5**). This indicates activation of the ATF6-CHOP-GRP78 axis in the ERAD response. We predict that this rapid response is to adapt to the cellular stress induced by injury and may induce repair. Interestingly, we also observed significant ubiquitination and enhanced gene expression levels of DDIT4, HSPa1B (**Figure 4A & data not shown**) post injury highlighting the involvement of a wider range of ER stress linked proteins to the injury cellular responses. However, we also observed that osteoarthritic tissues (cartilage or synovium) seem to fail to maintain this response and the ER stress genes are significantly reduced in diseased tissue compared to controls (**Supplementary Figure 1**). This highlights a possible disruption of ER homeostasis in OA, a mechanism that requires further investigation and potential druggable target development.

Our findings that injury rapidly caused a differential ubiquitination and activation of the ER associated deubiquitinases; YOD1 and ATXN3 indicated that this change in ubiquitination pattern might directly regulate protein activity. Many DUBs are associated with E3 ligases that have an intrinsic tendency to self-ubiquitinate. This interaction allows stabilisation of the E3 ligase by reversing its auto-ubiquitination and preventing its degradation. Likewise, DUBs are also targets for ubiquitination by E3 ligases [8–10, 44–48]. E3-DUB interactions may allow fine-tuning of the ubiquitination status of a common substrate [7, 46]. Ubiquitination of DUBs may regulate their catalytic activity by either competing with binding of ubiquitinated substrates especially when associated with an E3 ligase or by enhancing activity through activating conformational changes [8–10]. The ubiquitination of ATXN3 at Lysine 117 was reported to enhance its activity *in vitro* [8] and *in vivo* [49]. The ubiquitination of YOD1 at Lysine 213/257 is a novel observation that is not reported previously. We observed that YOD1 is mostly deubiquitinated after injury (**Figure 4C**) and this may have caused the enhanced activity of a shifted form of the protein as observed in **Figure 6B**. This high molecular weight form could be a dimer or multimer.

Interestingly we observed that YOD1 and ATXN3 are also active in osteoarthritic cartilage (**Figure 6E**), highlighting a possible sustained activation of these two DUBs following repetitive injury in diseased tissue. Inhibition of DUB activity using the broad-spectrum DUB inhibitor PR-619 in a zebrafish model of tissue injury caused a significant reduction in injury-induced inflammatory response as seen by reduced number of neutrophils recruited to wound site (**Figure 7**) and reduced injury-dependent NFkB activation (**Figure 8**). This implicates DUBs in modulation of inflammatory cellular responses to localised connective tissue injury. Further investigation of the role of these DUBs in modulating the pathological injury and disease progression *in vivo* is a crucial future prospective for validating these data *in vivo*. Another important question is if modulating DUB activity would interfere with cartilage degradation *in vivo* or *in vitro*. The role of these modified sites in cartilage development and function *in vivo* could be addressed in future mutational studies. DUBs are attractive therapeutic targets in several diseases and we expect in the near future a number of clinical trials would include the use of DUB inhibitors or drugs that showed anti-DUB properties in osteoarthritis patients. The identification of specific inhibitors for injury-induced DUBs is another promising new avenue in disease targeting.

Overall, in this study we identify a unique ubiquitin signature associated with injury cellular responses. A specific set of proteins incorporating deubiquitinases and ER stress regulators is also identified to be modulated by ubiquitination. We show that injury-dependent differential ubiquitination facilitated the activation of the 3 DUBs USP5, YOD1 and ATXN3. The early cellular responses to injury involves ER stress markers activation. Using data mining approaches, we observed that ER stress genes are significantly down regulated in human osteoarthritic cartilage while negative regulators of ER stress as HSPA8 were significantly upregulated, highlighting a disrupted ER homeostasis in OA. The work presented here provides a number of significant clues of the early cellular responses to tissue injury that could be modulated to delay or inhibit tissue damage in diseases induced by pathological injury such as osteoarthritis.

## 3. Materials and Methods

### 2.1. Cartilage *ex vivo* injury model and preparation of cartilage tissue lysates

Porcine articular cartilage was obtained from the metacarpophalangeal (MCP) joints of the pig forefeet or trotters. Pig trotters of freshly slaughtered 3–6-month-old pigs were purchased from a local farm. Trotters were first decontaminated in 2% Virkon, and then equilibrated at 37°C for 1hour before use. MCP joints were opened under sterile conditions and injury to cartilage was induced by explantation from the articular surface as previously described [5]. Cartilage was either snap-frozen (at time point 0) or cultured for indicated times in serum-free DMEM culture medium.

### 2.2. Preparation of cartilage tissue lysates and enrichment of ubiquitinated Gly-Gly peptides using immunoafinity purification

Cartilage tissues were explanted as described above from the articular surface of MCP joints of 4 pig trotters and pooled as one replicate for each time point. Cartilage tissues (4g) for each replicate were lysed in 10ml Urea Lysis buffer containing Urea (9M) HEPES buffer (20mM) pH 8.0, Sodium orthovanadate (1mM), Sodium pyrophosphate (2.5mM) and β-glycerophosphate (1mM) for 2hrs at room temperature with continuous vortex mixing.

Tissue lysates were then sonicated three times using 15W microtip for 15 seconds each. Samples were cooled on ice for 1 min between each sonication burst then cleared by centrifugation at 13000rpm for 15 minutes. Supernatants were transferred to new tube and then reduced with 1.25mM DTT for 30 minutes at 55°C. Extracts were then cooled down to room temperature then alkylated using 1.9mg/ml Iodoacetamide for 15 minutes in the dark. Proteins were then digested overnight with trypsin at room temperature. Peptides were then purified using Sep-Pak^®^ C18 columns. 50μg of tryptic peptides were saved for global proteome analysis. Gly-Gly peptides were then purified using Ubiquitin remnant motif (K-ε-GG) kit from Cell Signalling Technology (5562). Tryptic peptides were incubated with K-ε-GG immunoaffinity beads for 2hrs at 4°C then washed as per manufacturer’s instruction to remove unbound peptides. Gly-Gly enriched peptides were eluted using 0.15% TFA then concentrated and purified for LC-MS/MS analysis as per manufacture’s procedure.

### 2.3. Mass spectrometry raw data processing, visualisation and pathway analysis

Peptide material from pulldown experiments was re-suspended in 20μl water with 0.1% trifluoroacetic acid and 2% acetonitrile. Samples were then injected into a nano-UPLC U3000 system (Thermo Fisher) coupled to an orbitrap Fusion Lumos tandem mass spectrometer (Thermo Fisher) as described previously [50].

#### Raw Data, data visualisation and pathway analysis

Mass spectrometry raw data were processed using MaxQuant software package (version 1.6.7.0). Default settings were used. Variables modifications included diGly remnant of Lysine and Methionine oxidation/acetylation while fixed modification was set as carbamidomethylation. Protein searches were performed using Sus Scrofa/ Homo sapiens protein sequences uniprot fasta file. Processed data was analysed by Label Free Quantification (LFQ) and visualised using Perseus software package tools (v1.6.12.0)[51] and Venny 2.1 (https://bioinfogp.cnb.csic.es/tools/venny/). Downstream analysis of the ‘proteinGroups.txt’ and ‘GlyGly. Txt’ output tables for proteome and enriched ubiquitinated peptides respectively were performed in Perseus. Columns for experiment and control set were selected and log transformed. Quantitative profiles were filtered for missing values, and were then filtered independently for each of time point and control pairs, retaining only proteins and ubiquitin peptides that were quantified in all three replicates of either the injury time point or control pull-down. Missing values were imputed (width 0.3, down shift 1.8) before combining the tables and performing the multi-volcano analysis. Data quality and normal distribution was inspected and represented in a multi-scatter plot and histogram (**supplementary figures 2 & 3**). Pathways and functional enrichment analysis of enriched hits and proteinprotein interactions were performed using STRING v11.0 database [29]. Heat maps were generated using the online platform Morpheus/ Broad institute (https://software.broadinstitute.org/morpheus/).

### 2.4. Validation of enriched ubiquitinated proteins: Ubiquitin enrichment pull down assays and western blotting

Cartilage was snap frozen immediately at the end of time points. Proteins were extracted from cartilage by lysis in 2 volume of autoclaved glass beads lysis buffer (50mM Tris, 5mM MgCl2, 0.5mM EDTA, and 250mM sucrose) and 1 volume of acid washed glass beads (Sigma, G4649). 1mM DTT was added fresh to samples with vortex mixing at 4°C for 2-3 hrs. Samples were then pelleted, clear supernatants were collected, and protein concentration was quantified using Qubit 4 fluorimeter as per manufacture’s procedure (Invitrogen). Ubiquitinated proteins were enriched from cleared tissue lysates (500μg) using anti-ubiquitin antibody conjugated beads (anti-mouse, Santa Cruz) overnight at 4°C with rotation. Beads were pelleted then washed with lysis buffer for 5 times/5 min each. enriched proteins were then eluted by heating beads in 1X sample buffer. Further analysis of ubiquitinated proteins was performed by western blotting using protein specific antibodies. Protein A agarose beads were used as a negative control for non-specific pull down. Antibodies used in this analysis were from ProteinTech against VCP (10736-1-AP), Ubiquilin 1(23516-1-AP), UBE2N(10243-1-AP), UBE2L3(14415-1-AP), USP5 (10473-1-AP), RAD23B(12121-1-AP), YOD1(25370-1-AP), BRCC3(15391-1-AP), ATXN3(13505-1-AP), GRP78/BIP(11587-1-AP), DDIT3/CHOP(15204-1-AP) and β-actin (66009-1-Ig).

### 2.5. Deubiquitinases activity assays and active DUBs pull down

#### Using activity probes

TAMRA-Ub-VME (UbiQ-050) and HA-Ahx-Ahx-Ub-VME (UbiQ-035) were purchased from UbiQ Bio BV, NL. Porcine cartilage and human osteoarthritic cartilage tissues lysates were reacted with DUB activity probe for 45 min at 37°C. Lysates reacted with HA activity probe were then used for active DUBs pull down using Anti-HA conjugated magnetic Beads (88837, Thermo Scientific, UK). Immunocomplexes were then washed three times using lysis buffer then analysed by western blotting using DUB specific antibodies. Lysates reacted with TAMRA probe were analysed by SDS-PAGE followed by imaging fluorescence using Syngene gel imaging system.

#### Ubiquitin-rhodamine (110)-glycine assay

DUB activity in cartilage tissue lysates were measured using ubiquitin-rhodamine(110)-glycine quenched substrate (Boston Biochem, cat No U-555). In the presence of active DUBS, the amide bond between the ubiquitin C-terminal glycine and rhodamine110Gly results in an increase in fluorescence at 535 nm (Exc. 485 nm). 5-10 μg of tissue lysates were incubated with 250nM substrate and fluorescence intensity reflecting activity was measured over time at 10 min intervals using a Varioskan™ LUX multimode microplate reader (Thermo Scientific, UK). Readings were retrieved from ScanIT software and Z scores were calculated and plotted against time using GraphPad prism.

### 2.5. Zebrafish experiments

#### 2.6.1 Husbandry and maintenance

Zebrafish wildtype and transgenic lines adult fish were maintained on a 14:10-hour light/dark cycle at 28°C in The Bateson Centre aquaria (University of Sheffield), under Animal Welfare and Ethical Review Bodies (AWERB) and UK Home Office-approved project license protocols.

#### 2.6.2 Tail fin injury assays

Tail fin injury was induced in zebrafish embryos by tail transection as described previously [38]. Zebrafish larvae were anesthetized at 3 or 5 dpf by immersion in tricaine (0.168 mg/ml), and tail fin injury was induced by fin fold/tail transection using a sterile microscalpel. Embryos were then incubated in E3 medium at 28°C as per indicated times in each experiment. Following incubation embryos were either mounted in 1% low melting agarose for live imaging or fixed for in situ hybridisation or immunofluorescence.

#### 2.6.3 Inflammation assay

Recruitment of neutrophils to injury site in zebrafish embryos is a well-established assay for measuring inflammation post injury. For that, Tg(mpx:*EGFP*)*i114* zebrafish transgenic line was used to study neutrophils recruitment post injury. Zebrafish embryos at 3dpf were incubated with either DMSO or PR-619 DUB inhibitor (5-50μM) for 6 hrs or overnight before injury. Neutrophils were then counted at the site of transection at 2 hours post injury (hpi) using a fluorescence dissecting stereomicroscope (Leica).

### 2.6.4 Whole mount *in situ* hybridisation

Zebrafish GRP78/BIP and IL1 RNA probes were generated using PCR based method then DIG-labelled as described before[52]. For whole mount in situ hybridisation, larvae were fixed in 4% methanol-free Paraformaldehyde (PFA) in PBST(0.1% Tween20 in 1xPBS) for 24hrs at 4°C with gentle rocking then dehydrated in a methanol-series (25%-100%) in PBS and stored at −20°C overnight. Larvae were then rehydrated then washed in PBST for 4 times (5 minutes each). Samples were then digested with 10 μg/ml Proteinase K in PBST(Invitrogen,, USA). Larvae were then washed briefly in PBSTthen re-fixed for 20 min in 4% PFA-PBST followed by washes in PBST then incubated at 67 °C for 2 h in pre-warmed hybridisation buffer. Hybridisation buffer was replaced with probe buffer containing DIG labelled RNA probes for either IL1 or GRP78 diluted in hybridisation buffer and incubated at 67 °C overnight. Larvae were washed thoroughly at 67 °C with hybridisation buffer then washed in PBST and incubated for 1 h in blocking buffer with gentle agitation. Larvae were incubated overnight at 4 °C with anti-DIG antibody in blocking buffer then washed in PBST, followed by washes in staining wash solution. Larvae were then incubated in staining solution for 1hr and monitored for colour development. Reactions were stopped with stop solution when appropriate colour was achieved. Larvae were then fixed and dehydrated in methanol series as above, rehydrated then cleared with a series of glycerol dilution/H20/0.1%Tween20. Stained larvae were stored in 75%glycerol:25%H20.0.1%Tween20 at −20°C until ready for imaging.

### 2.6.5 Whole mount immunofluorescence and confocal microscopy

Zebrafish Larvae (3-5dpf) were fixed in 4% Paraformaldehyde overnight at 4°C. Fixed larvae were then washed 3 times in PBSTX buffer (0.1% triton X100 in 1xPBS) then dehydrated in 100% methanol overnight at −20°C. Following washing once in 1x PBSTX, larvae were then incubated in blocking buffer (PBSTX + 0.2% BSA + 5% serum) for 2 hours at room temperature with gentle rocking. Larvae were then incubated in primary antibodies (CHOP or GRP78) overnight at 4°C in blocking solution at 1:300 dilutions. Antibody solution was then removed and larvae was extensively washed with PBSTX at least 6 times for 10 min each before incubation with Alexa Flour 647 conjugated goat-anti rabbit secondary antibody overnight at 4 °C at 1:300 dilutions in blocking buffer. Samples were then washed 6 times for 10 minutes each in PBSTX. Larvae were then transferred to imaging slides and arrayed as needed and tails were cut to facilitate mounting using SlowFade™ Glass Soft-set Antifade Mountant with DAPI (Invitrogen, Catalogue number S36920).

Images of the tail regions were captured using the Nikon A1 confocal microscope (20X). Images were analysed using FIJI software (NIH) using TrackMate Plugin.

### 2.6.6. Live imaging/timelapse of NFkB reporter zebrafish transgenic line

Zebrafish Larvae were treated with DMSO or PR-619 prior to injury induction. Following injury, larvae were mounted in 1% low melting agarose containing tricaine (0.168 mg/ml) in E3 medium. GFP signal at the wounding site was then detected using Spinning disc confocal microscopy and z stack images were collected every 30 minutes for 18 hours. Z stacks were then converted by maximum intensity projection and time-lapse images were analysed and GFP signal was quantified at the tail fin site using FIJI Software(NIH).

### 2.6. Human tissue samples and ethical approval

Human osteoarthritic cartilage samples were obtained directly after knee replacement surgery from the South Yorkshire and North Derbyshire Musculoskeletal Biobank (SYNDMB). Samples were collected with informed donor consent in full compliance with national and institutional ethical requirements (Sheffield REC 20/SC/0144), the United Kingdom Human Tissue Act, and the Declaration of Helsinki. Freshly collected samples were snap frozen immediately after surgery. Proteins were isolated from cartilage as described above then lysates used for DUB activity and pulldown assays as described in section 2.5.

### 2.8 RNA extraction and RT-qPCR

RNA was isolated from articular cartilage as described previously[2]. Briefly, cartilage tissues were homogenised in 1ml Trizol (Invitrogen) for 1 min on ice and the homogenate was left at room temperature for 5 minutes then mixed with 200 μl of BCIP solution. Samples were vortexed for 15 seconds then incubated again at RT for 5 min followed by centrifugation at 13000 rpm for 15 min at 4°C. Aqueous layer was then separated and mixed with equal volume of 90% ethanol then transferred to zymogen columns. RNA was bound to column filter followed by washing as per manufacture’s manual. DNA was digested on column using DNAse1 as per manufacture’s manual then RNA was eluted in 20μl ddH2O. Equal concentration of RNA (1μg) were used for reverse transcription reactions using superscript II reverse transcriptase for 2hrs at 37°C. cDNA was then diluted and used in RT-PCR reaction using cyber green master mix (Applied Biosystems). RT-PCR reactions were performed in QuantStudio Flex 7 instrument and data were analysed using Max studio quantification. CT values were used for calculations of ddCT and used for fold of change in gene expression compared to control and using β-actin as housekeeping gene. Primers sequence are provided in **Supplementary Table 1**.

## Declarations

### Ethics approval and consent to participate

Zebrafish wildtype and transgenic lines adult fish were maintained on a 14:10-hour light/dark cycle at 28°C in The Bateson Centre aquaria (University of Sheffield), under Animal Welfare and Ethical Review Bodies (AWERB) and UK Home Office–approved project license protocols. Human osteoarthritic cartilage samples were obtained directly after knee replacement surgery from the South Yorkshire and North Derbyshire Musculoskeletal Biobank (SYNDMB). Samples were collected with informed donor consent in full compliance with national and institutional ethical requirements (Sheffield REC 20/SC/0144), the United Kingdom Human Tissue Act, and the Declaration of Helsinki.

### Availability of data and materials

The mass spectrometry proteomics data have been deposited to the ProteomeXchange Consortium via the PRIDE [53] partner repository with the dataset identifier PXD030817. Any further data generated or analysed during this study are included in this published article [and its supplementary information files]. The datasets analysed for this study can be found in gene expression omnibus (GEO) under accession numbers GSE114007, GDS2126, GDS5401, GDS5403 and GSE155892.

### Competing interests

The authors declare that they have no competing interests.

### Funding

Work in this study was funded by Versus Arthritis Fellowship to HI, grant number 22048.

### Authors’ contributions

study design and conception (HI), funding acquisition(HI), Data acquisition and analysis (HI, APF, NK), Validation (HI, NK), Writing original draft(HI), Writing-review and editing (HI, MW, BK, NK, APF), Resources (HI, BK, MW)

---

**Supplementary Figure S1:**
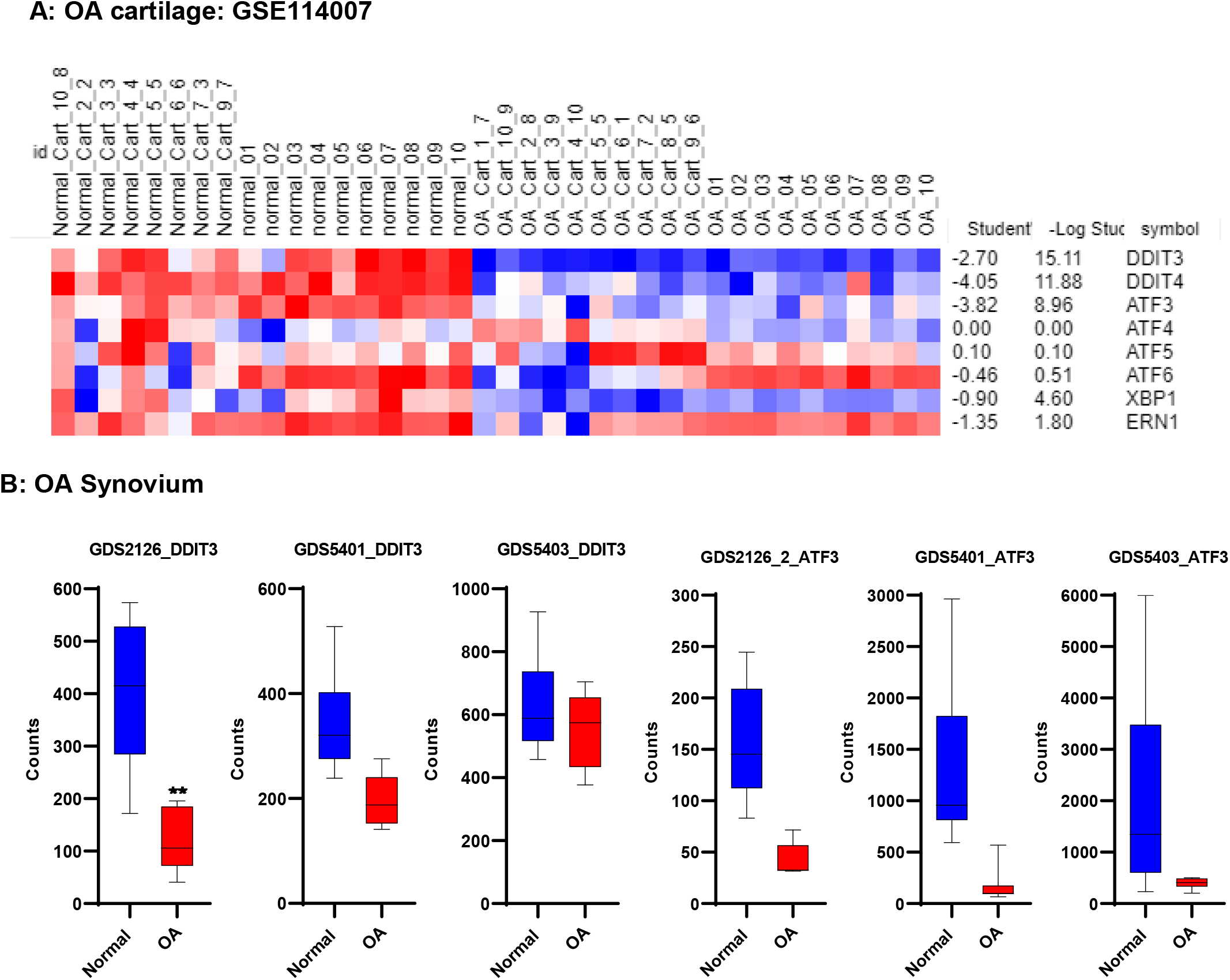
Modulation of ER stress responsive genes in osteoarthritic cartilage and synovium tissues compared to normal tissues. RNA counts were retrieved from Gene Expression Omnibus studies as following **GSE114007 RNA seq study in OA cartilage** versus normal donors (38 donors: 18 normal and 20 OA)(PMID: 30081074), GDS5403: 33 samples: 10 normal, 10 OA, 13 RA(PMID: 24690414) and GDS5401: Berlin data set (10 normal, 10 OA, and 10 RA)(PMID: 24690414)

**Figure 6-Supplement Figure 1:**
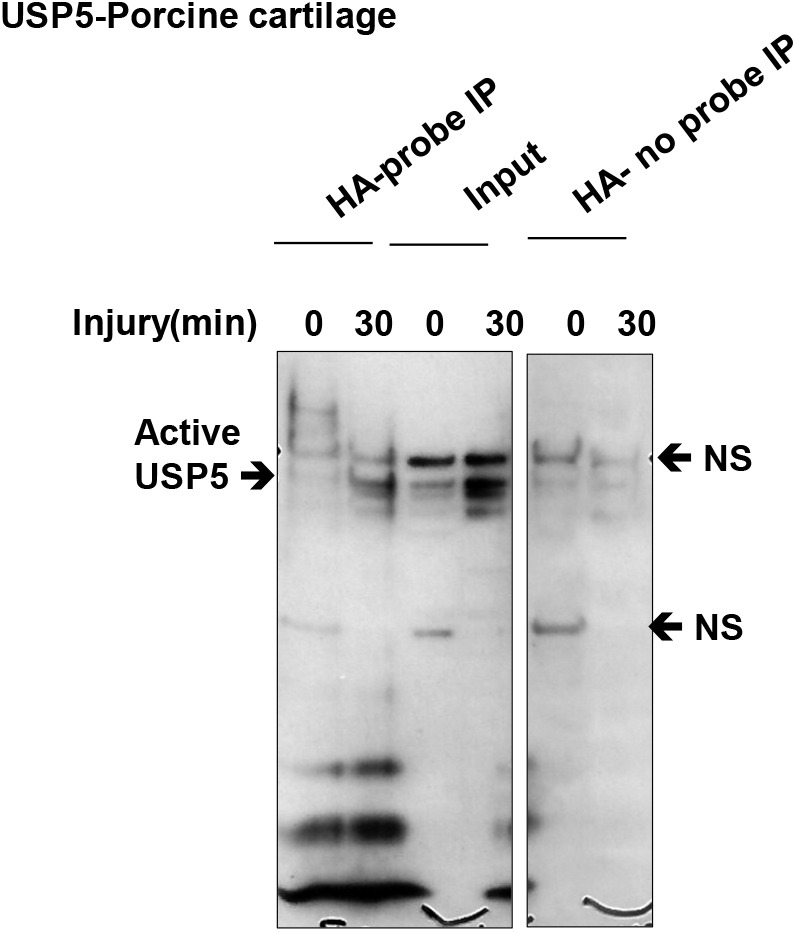
USP5 deubiquitinase is activated in response to articular cartilage injury. **Porcine cartilage was** lysed as described in methods. 50 μg of lysates were reacted with HA DUB activity probe and for 45 minutes at 37°C and then subjected to immunoprecipitation of active DUBs using anti HA antibody conjugated magnetic beads. Immunocomplexes were analysed by western blotting and compared with input and IP control using USP5 DUB specific antibody. NS: non specific

**Figure 7-Supplement Figure 1:**
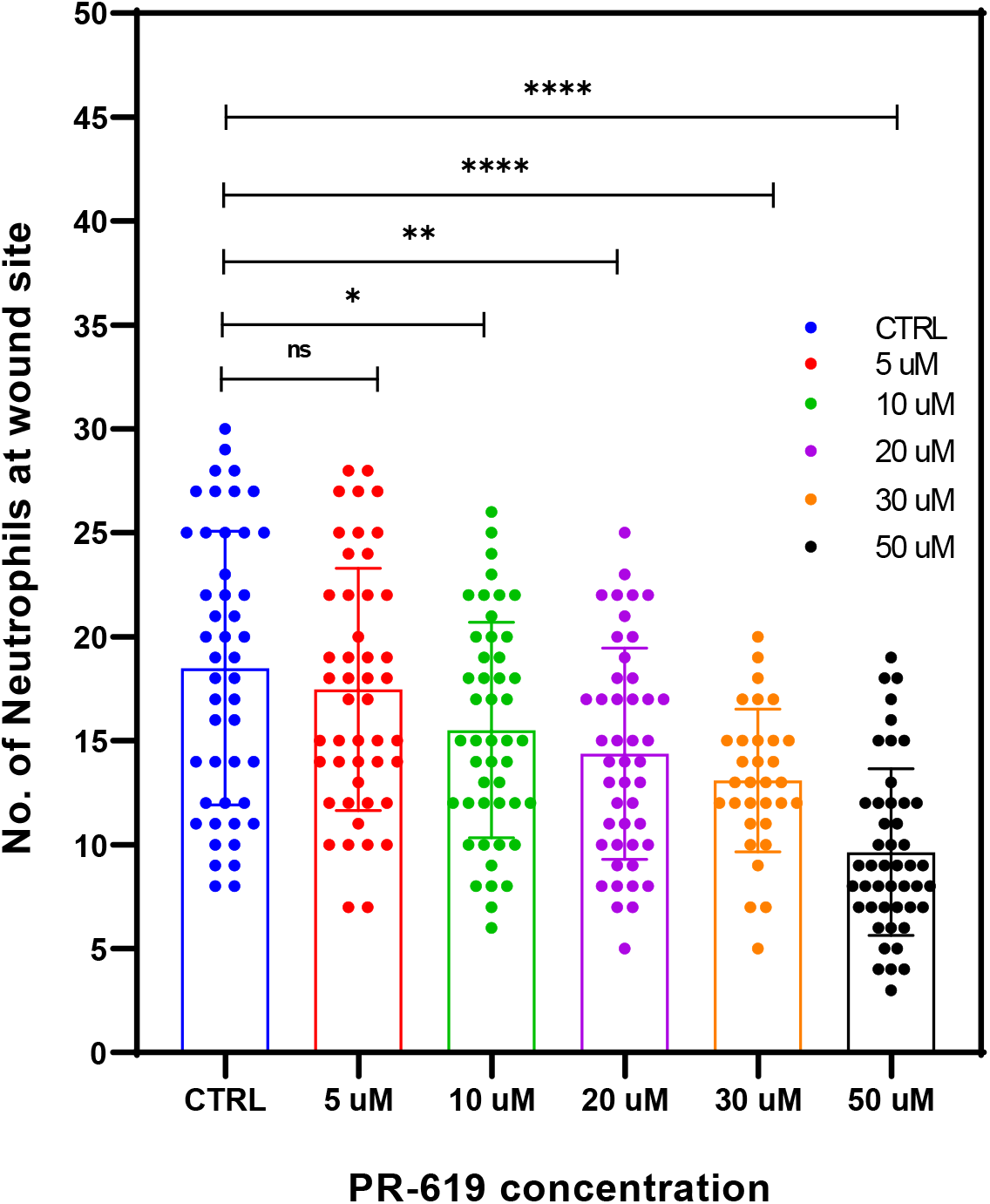
Effect of DUBs inhibition using PR-619 on neutrophil recruitment at wound site following tail fin injury in 3dpf Zebrafish Larvae. Tg(mpx:GFP)i114 neutrophil reporter line larvae(3dpf) were pre-treated with either DMSO as control or 5-50 μM PR-619 inhibitor for 6 hrs before tail fin injury. Neutrophils were then counted at the wound site 2hrs following injury. Statistics by unpaired t-test.

**Supplementary Figure 2:**
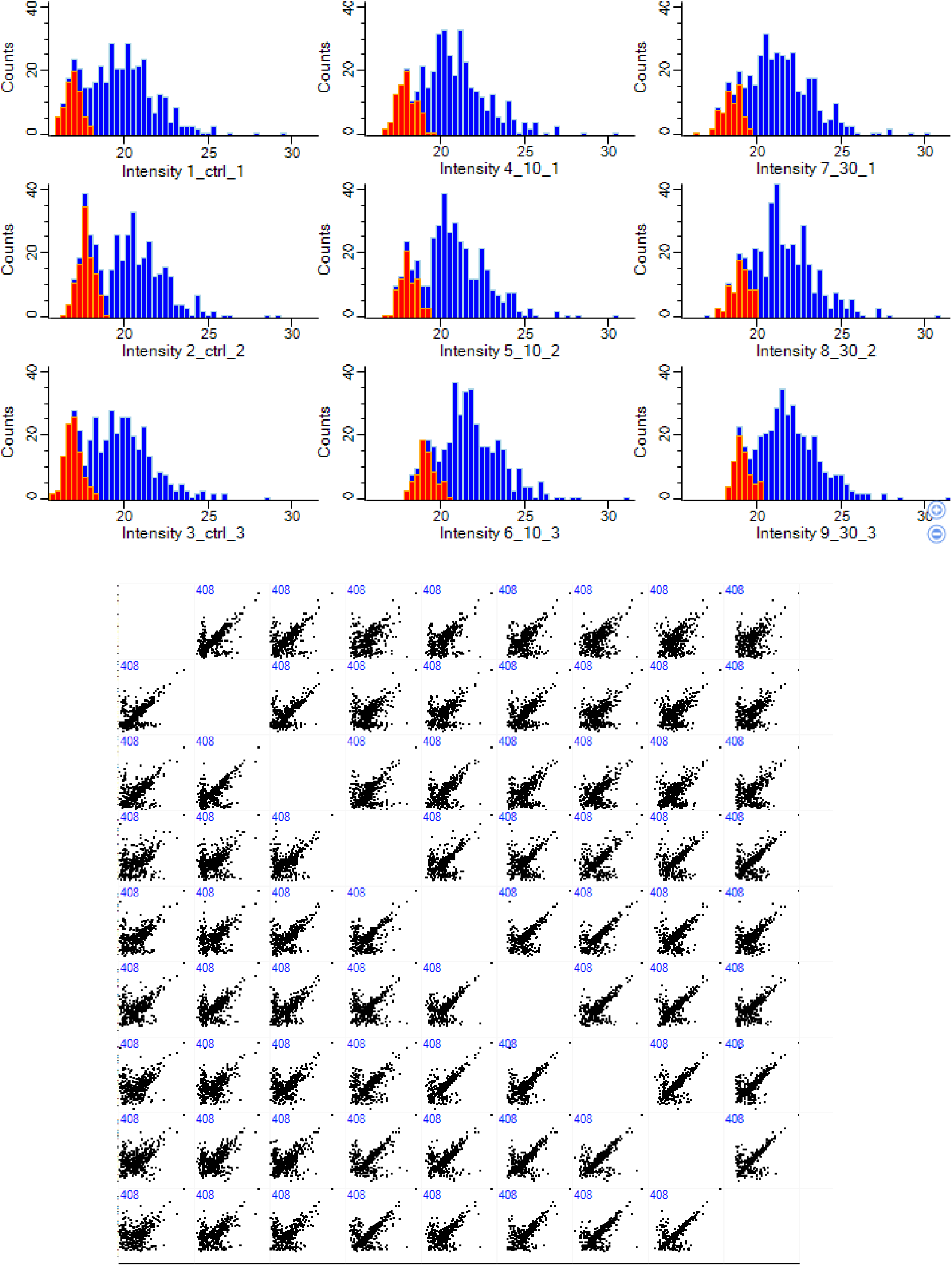
Histogram analysis and scatter plot of individual cartilage samples in ubiquitinome analysis. Missing values from normal distribution were imputed as described in methods(red color). Numbers on Scatter plot shows peptides of valid values in all samples(408)

**Supplementary Figure 3:**
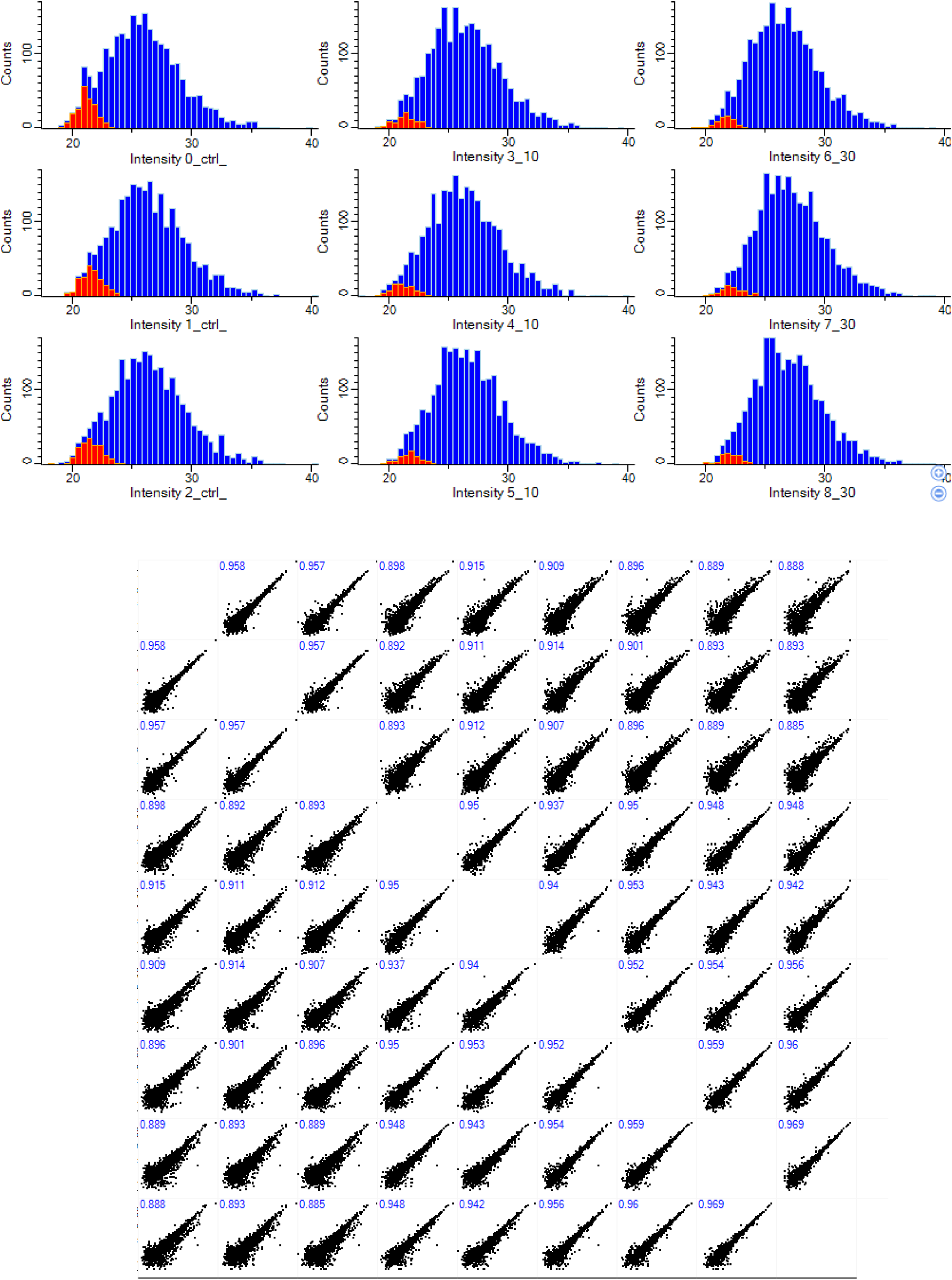
Histogram analysis and scatter plot of individual samples in proteome dataset. Missing values from normal distribution were imputed as described in methods(red color). Numbers on Scatter plot shows Pearson correlation between individual samples.

## Notes

### Competing Interest Statement

The authors have declared no competing interest.

